# Single-step genome-wide association study for resistance to *Piscirickettsia salmonis* in rainbow trout (*Oncorhynchus mykiss*)

**DOI:** 10.1101/587535

**Authors:** Rodrigo Marín-Nahuelpi, Agustín Barría, Pablo Cáceres, María E. López, Liane N. Bassini, Jean P. Lhorente, José M. Yáñez

## Abstract

One of the main pathogens affecting rainbow trout (*Oncorhynchus mykiss*) farming is the facultative intracellular bacteria *Piscirickettsia salmonis*. Current treatments, such as antibiotics and vaccines, have not had the expected effectiveness in field conditions. Genetic improvement by means of selection for resistance is proposed as a viable alternative for control. Genomic information can be used to identify the genomic regions associated with resistance and enhance the genetic evaluation methods to speed up the genetic improvement for the trait. The objectives of this study were to i) identify the genomic regions associated with resistance to *P. salmonis*; and ii) identify candidate genes associated with the trait. We experimentally challenged 2,130 rainbow trout with *P. salmonis* and genotyped them with a 57 K SNP array. Resistance to *P. salmonis* was defined as time to death (TD) and as binary survival (BS). Significant heritabilities were estimated for TD and BS (0.48 ± 0.04 and 0.34 ± 0.04, respectively). A total of 2,047 fish and 26,068 SNPs passed quality control for samples and genotypes. Using a single-step genome wide association analysis (ssGWAS) we identified four genomic regions explaining over 1% of the genetic variance for TD and three for BS. Interestingly, the same genomic region located on *Omy27* was found to explain the highest proportion of genetic variance for both traits (2.4 and 1.5% for TD and BS, respectively). The identified SNP in this region is located within an exon of a gene related with actin cytoskeletal organization, a protein exploited by *P. salmonis* during infection. Other important candidate genes identified are related with innate immune response and oxidative stress. The moderate heritability values estimated in the present study show it is possible to improve resistance to *P. salmonis* through artificial selection in the current rainbow trout population. Furthermore, our results suggest a polygenic genetic architecture and provide novel insights into the candidate genes underpinning resistance to *P. salmonis* in *O. mykiss*.

## INTRODUCTION

As in any intensive animal production system, infectious diseases are one of the main threats affecting the success and sustainability of aquaculture (Yáñez *et al.* 2014a). In the case of salmonid production, one of the major pathogens affecting productivity is the facultative intracellular bacteria *Pisciricketssia salmonis*, etiological agent of salmonid rickettsial syndrome (SRS). This bacterium was first identified in 1989 in Chile, in a farmed coho salmon (*Oncorhynchus kisutch*) population (Cvitanich *et al.* 1991). Since then, mortalities resulting from SRS have been also identified in Atlantic salmon (*Salmo salar)* and rainbow trout (*Oncorhynchus mykiss*) in several countries, such as Scotland, Ireland, Norway and Chile (Fryer and Hedrick 2003). In Chile, SRS was responsible for 20.7, 67.9 and 92.6% of the mortalities associated with infectious diseases in *S. salar*, *O. kisutch* and *O. mykiss*, species respectively (Sernapesca 2018). To date, strategies for *P. salmonis* control and treatment are mainly based on vaccines and antibiotics. The effectiveness of both approaches has not been adequate (Rozas and Enríquez 2014). Therefore, it has been estimated that economic losses due SRS mortalities, reached up to US$450 million in Chile in 2012 (Camussetti *et al.* 2015). However, variables such as laboratory diagnosis screening expenses or loss of quality of the harvested fish and products were not considered, implying that the economic impact could be even higher.

Therefore, selective breeding could be a feasible alternative to enhance disease resistance; reducing mortality rates from *P. salmonis*, as well as improving animal health and productivity (Bishop and Woolliams 2014: Yáñez and Martínez 2010). However, the main requisite to include a trait into a genetic program is the presence of significant additive genetic variance within the population (Falconer and Mackay 1996). Previous studies estimated heritability values ranging from 0.11 to 0.41 for *P. salmonis* resistance in Atlantic salmon and coho salmon (Yáñez *et al.* 2013; Yáñez *et al.* 2016a; Barría *et al.* 2018). In the case of rainbow trout, Yoshida *et al.* (2018a) estimated heritabilities ranging from 0.39 to 0.57 for resistance to *P. salmonis* using day of death and 0.54 to 0.62 for binary survival as trait definitions. Altogether, these results demonstrate the possibility of improving this trait by means of artificial selection in different salmonid species.

The development of next generation sequencing technologies has facilitated the identification of thousands of single nucleotide polymorphisms (SNPs) segregating along the genome of several animals, including aquaculture species (Yáñez *et al.* 2015). Thus, using a genotyping by sequencing (GBS) approach in conjunction with genome-wide association studies, some authors evaluated genomic regions associated with resistance to bacterial infections in aquaculture species (Liu *et al.* 2015; Palti *et al.* 2015a; Palaiokostas *et al.* 2016; Barría *et al.* 2018). However, in salmonid species, the use of SNP panels has been the most used alternative for genotyping a high number of individuals with thousands of genetic variants simultaneously. This has been made simpler by the development of high density SNP arrays for Atlantic salmon (Houston *et al.* 2014; Yáñez *et al.* 2016b) and rainbow trout (Palti *et al.* 2015b). The use of these SNP panels have also allowed the comparison of the accuracy of estimated breeding values (EBV) using genomic selection to pedigree-based genetic evaluations for resistance to infectious diseases in Atlantic salmon (Ødegård *et al.* 2014; Tsai *et al.* 2016; Bangera *et al.* 2017; Correa *et al.* 2017), coho salmon (Barría *et al.* 2018) and rainbow trout (Vallejo *et al.* 2016; Vallejo *et al.* 2017a; Yoshida *et al.* 2018a; 2018b). SNP arrays have also enabled the dissection of the genetic architecture of resistance to bacterial diseases in salmonids. For instance, genomic regions and candidate genes associated with resistance to *P. salmonis* in Atlantic and coho salmon (Correa *et al.* 2015; Barría *et al.* 2018), and bacterial cold water disease (BCWD) in rainbow trout (Vallejo *et al.* 2017b) have been identified.

To date there are no studies aimed at identifying genomic regions or candidate genes associated with resistance to *P. salmonis* in rainbow trout populations. Therefore, the main objectives of the current study were to i) identify genomic regions associated with resistance to *P. salmonis* in a farmed rainbow trout population, and ii) identify candidate genes associated with the trait.

## MATERIALS AND METHODS

### Population and experimental challenge

The population used in this study was rainbow trout (*Oncorhynchus mykiss*) year-class 2011 bloodstock, owned by Aguas Claras (Puerto Montt, Chile) and belonging to a genetic improvement program run by Aquainnovo (Puerto Montt, Chile). This population was artificially selected for growth, appearance-related traits and carcass quality for three generations. For a detailed description about rearing conditions and population management please see Flores-Mara *et al.* (2017), Rodríguez *et al.* (2018) and Neto *et al.* (2019)

Fish from 105 full-sib families (48 half-sib families) with an average weight of 7.0 ± 1.5 g, were PIT-tagged for individual traceability of families. After tagging, fish were maintained in a single tank until they were transferred to Aquainnovo’s Aquaculture Technology Center Patagonia in August 2012. Fish were acclimated for 20 days in a 15m^3^ tank, prior to experimental challenge. A random sample of fish were selected to evaluate the sanitary status of the population, i.e. qRT-PCR for Infectious Salmon Anemia virus (ISAV), Infectious Pancreatic Necrosis virus (IPNV), and *Renibacterium salmoninarum*, and culture for *Flavobacterium spp*. Later, a total of 2,130 juveniles (with an average of 23 individuals per family and ranging from 17 to 27 fish per family), were intraperitoneally (IP) injected with 0.2ml of a lethal dose (LD_50_) of the LF-89 strain of *P. salmonis* inoculum. Post injection, fish were equally distributed into three different tanks, considering similar family distribution into each replicate (with 5 to 9 fish per family in each tank). Environmental parameters were measured throughout the challenge and the experimental challenge continued until the mortality curve showed a plateau. Daily mortality was recorded, and body weight was measured for each fish at time of death or at the end of the experiment (FW). Surviving fish were euthanized and body weight was also recorded. Fin clips from all fish were sampled and stored in 95% ethanol at −80°C until they were genotyped.

### Genotyping

The genomic DNA from the sampled fin clips was extracted using a commercial kit (DNeasy Blood & Tissue Kit, Qiagen), following the manufacturer’s instructions. Genotyping was performed using a commercial 57K SNP array (Affymetrix® Axiom® myDesignTM SNP) developed by the National Center for Cool and Cold water Aquaculture at USDA (Palti *et al.* 2015b).

Quality control (QC) was assessed through Affymetrix’s Axiom Analysis Software, using default settings. Then, a second QC using Plink software (Purcell *et al.* 2007), was applied to remove SNPs with a genotype call rate lower than 0.90, minor allele frequency (MAF) < 0.01 and deviated from Hardy-Weinberg Equilibrium (p < 1×10^−6^). Individuals with a call rate lower than 0.90 were also removed from further analyses.

### Trait definition

Resistance to *P. salmonis* was defined as time to death (TD), measured in days, with values ranging from 1 until the end of challenge test. Additionally, resistance to *P. salmonis* was also defined as binary survival (BS), with a value of 1 or 0 based on if the fish died or survived until the end of the challenge.

### Genomic-Wide association study

A single-step GWAS (ssGWAS) analysis was performed to identify genomic regions associated with resistance to *P. salmonis*. This approach considered fish with both phenotypes and genotypes and also individuals with phenotypes but no genotypes in the analysis (Wang *et al.* 2012). The pedigree and genotypic data in ssGWAS are connected through the H matrix. Thus, the H matrix combines both the pedigree and the genomic relationship matrices (Aguilar *et al.* 2010). Thus, the inverse of the H matrix is:

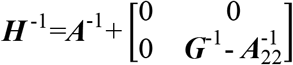

Where *A*^−1^ is the inverse of the numerator relationship matrix, considering all the phenotyped animals, 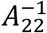 is the inverse of the pedigree-based relationship matrix considering only the genotyped animals, and *G*^−1^ is the inverse genomic relationship matrix. The following model was used for GWAS analysis:

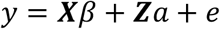

Where *y* is the vector of phenotypes (for TS and BS), *β* is the vector of fixed effects (tank as factor and final body weight as a covariate), *a* is the vector of random effects, *e* is the vector of residuals, and ***X*** and ***Z*** are the incidence matrices for fixed and random effects, respectively. A linear model and a threshold model were used for TD and BS, respectively. Both trait definitions were fitted using BLUPF90 statistical software (Misztal *et al.* 2016). Thus, AIREML and THRGIBBS1F90 were used for TD and BS, respectively. For the latter, a total of 200,000 Markov Chain Monte Carlo (MCMC) iterations were used, the first 20,000 were discarded as burn-in iterations and from the remaining 180,000 samples, we saved one from every 50. Therefore, the analyses included 3,600 independent samples.

For TS and BS, the heritability was estimated as follows:

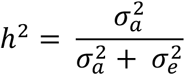

Where 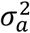 is the additive genetic variance estimated using the H matrix, and 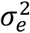 is the residual variance.

To identify genomic regions associated with each trait, we estimated the percentage of the genetic variance (PGV) explained by windows of 20 adjacent SNPs. Then, if a 20 SNP window explained more than 1% of the PGV, we considered that region as associated with resistance to *P. salmonis*.

### Candidate genes

The candidate genes were identified by searching 500kb up and downstream from the SNP explaining the highest proportion of PGV within each associated window. For this purpose, we used the last version of the *Oncorhynchus mykiss* reference genome (GCA_002163495.1). The criteria for selecting candidate genes lies in the function of the protein that encodes each gene found, mainly related to immune response, DNA repair, stress response and similar pathways.

### Data availability

The raw genotypes and phenotype data are available from the online repository figshare (https://figshare.com/s/221a39319b236d46f9fc). Table S1 contains all genes located within 1Mb window surrounding the SNPs explaining the highest proportion of genetic variance and is available at 10.6084/m9.figshare.7883342.

## RESULTS

### Descriptive statistics and heritabilities

Summary statistics for resistance to *P. salmonis* measured as TD and as BS and for FW are shown in Table 1. The first death was recorded on day 10 post intraperitoneal injection; the last on day 32. Average TD was 23.26 ± 7.86 days. At the end of the experimental challenge the proportion of non-survivor fish was 0.59 ± 0.49. Cumulative mortality among all 105 families ranged from 7.7 to 100%, indicating considerable phenotypic variation for resistance to *P. salmonis* in the current rainbow trout population. Cumulative mortality within each replicate tank was 59.4, 65.1 and 64.7%. Mortality peaked on days 12, 15 and 19 post injection. Average final body weight was 173.80 ± 52.27 g. This trait ranged considerably among challenged fish, with a minimum of 46.10g and maximum 448g.

**Table 1.** Summary statistics for time to death (TD), binary survival (BS) and final weight (FW) measured in 2,130 rainbow trout individuals.

| Trait | Mean | SD | CV(%) | Min | Max |
| --- | --- | --- | --- | --- | --- |
| TD | 23.26 | 7.86 | 33.27 | 10 | 32 |
| BS | 0.59 | 0.49 | 0.83 | 0 | 1 |
| FW | 173.80 | 52.27 | 30.07 | 46.10 | 448 |

Variance components for TD and BS are shown in Table 2. Significant heritability values were estimated for both trait definitions. Thus, 0.48 ± 0.04 and 0.34 ± 0.04 were estimated for TD and BS, respectively. Furthermore, a high genetic correlation was found between both traits (−0.96 ± 0.01).

**Table 2.**
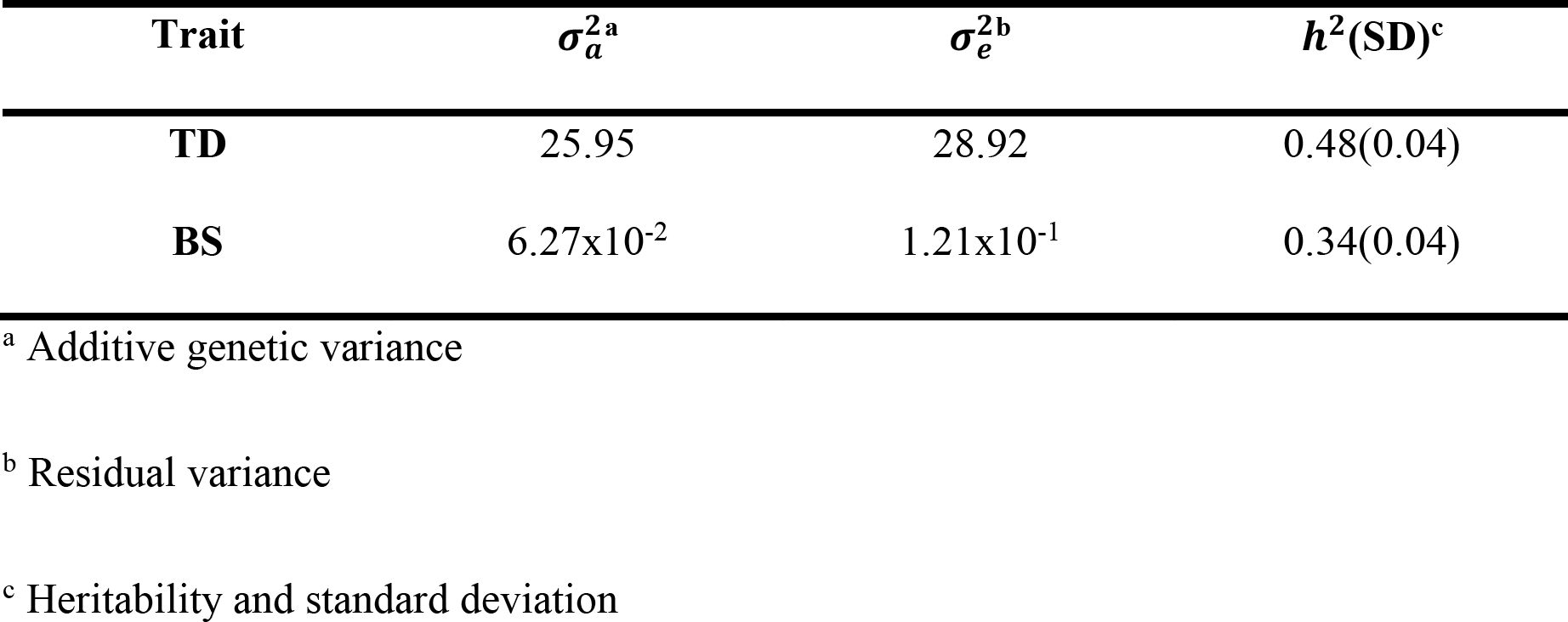
Genetic parameters and heritabilities for resistance to *Piscirickettsia salmonis* as time to death (TD) and binary survival (BS).

| Trait | $\sigma_a^2$<sup>a</sup> | $\sigma_e^2$<sup>b</sup> | $h^2$(SD)<sup>c</sup> |
| --- | --- | --- | --- |
| TD | 25.95 | 28.92 | 0.48(0.04) |
| BS | 6.27x10<sup>-2</sup> | 1.21x10<sup>-1</sup> | 0.34(0.04) |

<sup>a</sup> Additive genetic variance
<sup>b</sup> Residual variance
<sup>c</sup> Heritability and standard deviation

### Genome-wide association study

From all genotyped animals, 2,047 passed quality control (representing 97.10% of the total). A total of 26,068 SNPs remained in the set for further analyses (~ 64.68%). The Figure 1 shows the Manhattan plot for resistance to *P. salmonis* measured as TD and BS. We identified four genomic regions associated with resistance as TD. These regions were located on *Omy3*, *Omy14*, *Omy24* and *Omy27*. For BS, we identified three genomic regions associated with the trait. These were found on *Omy5*, *Omy27* and *Omy30*. Interestingly, the genomic region located on *Omy27* was found to be associated with resistance to *P. salmonis* for both TD and BS. In both cases, this common genomic region explains the highest proportion of genetic variance for each trait, with 2.4 and 1.5% for TD and BS, respectively. The SNP explaining the highest proportion of the genetic variance (Affx-88923370) is the same for both TD and BS (Table 3).

**Table 3.**
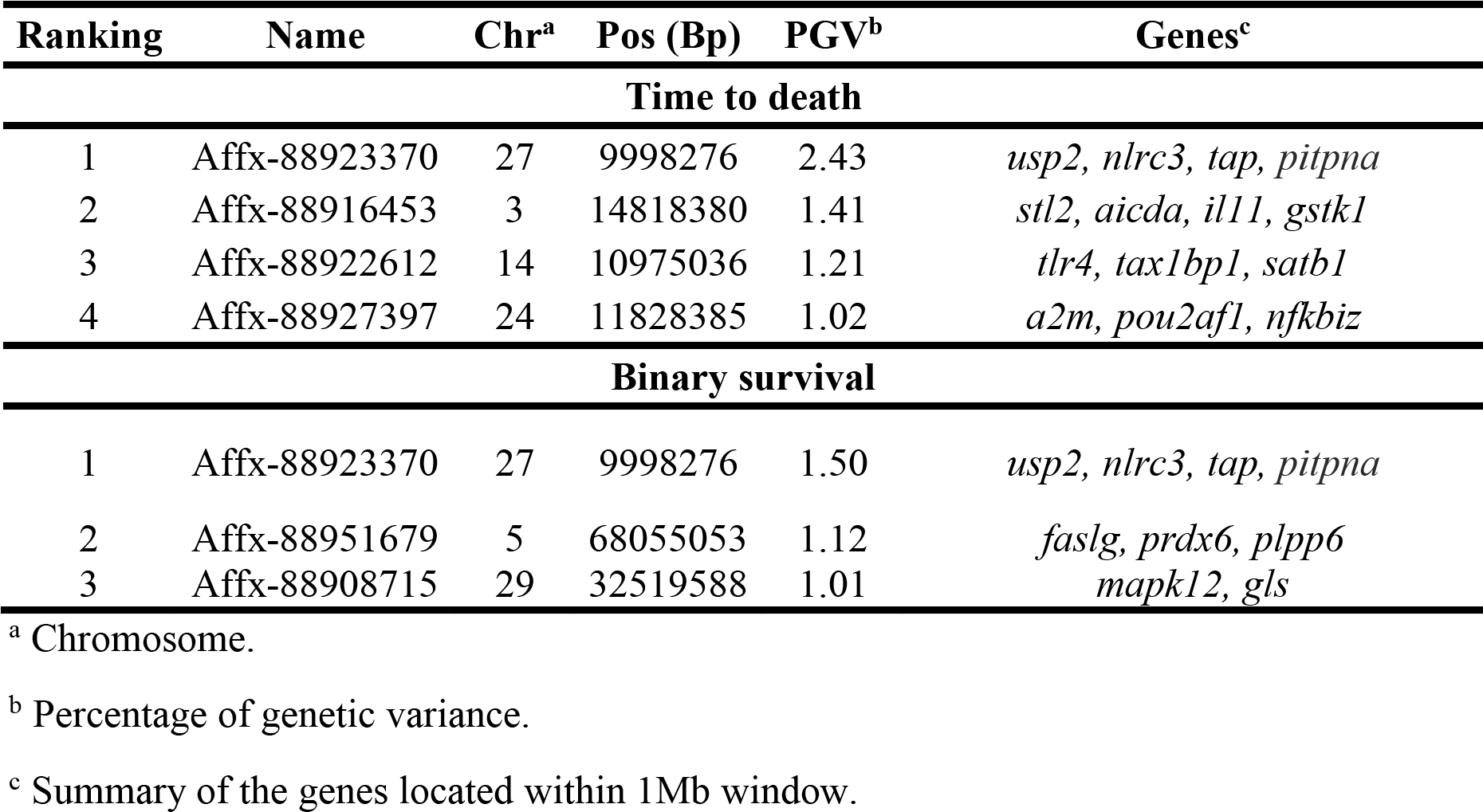
Top markers associated with *Piscirickettsia salmonis* resistance defined as TD and BS in rainbow trout, using ssGWAS,

**Figure 1.**
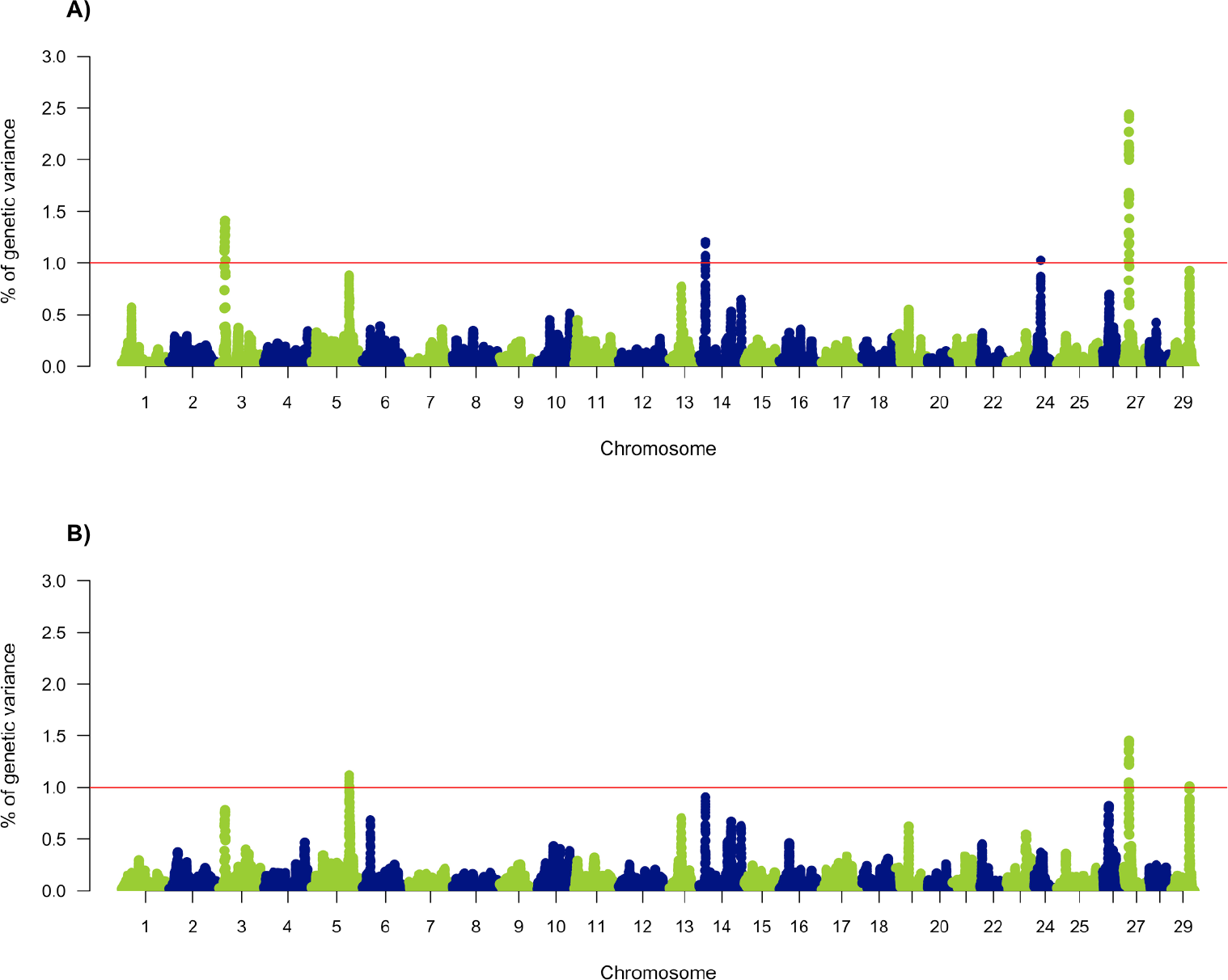
Genomic association analysis for resistance to *Piscirickettsia salmonis* in rainbow trout (*Oncorhynchus mykiss*). Resistance was defined as time to death (A) and as binary survival (B).

Using the *O. mykiss* reference genome (GCA_002163495) we identified candidate genes associated with resistance to *P. salmonis*. Table 3 shows a summary of the genes located proximate to the SNPs explaining the highest proportion of the genetic variance within each genomic region.

Among the candidate genes flanking the most important SNP on *Omy3* for TD, we found *Gluthatione S-transferease kappa 1* (*gstk1*) and Interleukin-11 (*il11*). These genes are involved in the response to oxidative stress and the immune response to bacterial infections, respectively (Oruc *et al*. 2004; Wang *et al*. 2005). On *Omy14*, we found the *Toll-like receptor 4* (*tlr4*) gene, which has been suggested to act as a bacteria sensor (Palti 2011). On *Omy24*, we found *alpha-2-macroglobulin-like* (*a2m*), which is part of a broad-spectrum protease inhibitor, and it has been suggested that plays a role in the defense against *Cryptobia salmositica* on rainbow trout (Zuo and Woo 1997).

Also, we found *POU class 2 associating factor 1* (*pou2af1*), which has been described as a coactivator of transcription factors that regulate Ig expression of B cells (Teitell 2003), and *NF-kappa-B inhibitor zeta-like (nfkbiz)*, a regulator of process like the pathogen recognition, phagocytosis and production of cytokines by dendritic cells (Rozas-Serri *et al.* 2017).

For BS, on *Omy5* we found *fas ligand* (*faslg*), whose protein has been suggested as an important mediator of anti-bacterial innate immune response, by inducing apoptosis of target cells and recruiting phagocytic cells (Kaur *et al.* 2004). On the same chromosome we found *Peroxiredoxin-6-like* (*prdx6*), one of the six different isoforms that conforms the peroxiredoxins group, which are antioxidants proteins that protect cells from oxidative damage and is likely to be involved in protective response against a bacterial infection in *Scophthalmus maximus* (Zheng *et al*. 2010).

On *Omy29*, *MAPK12* was found; previous studies described that MAPK12 is involved on the signaling pathways responsible for TNF-α secretion from rainbow trout macrophages, there for in innate immunity (Roher *et al*. 2011). *Glutaminase kidney isoform, mitochondrial-like* (*gls*) was also found on *Omy29*, which family proteins, generally forms a part of enzymes that plays a role in nucleotide, amino acid and urea biosynthesis (Kumada *et al.* 1993).

On *Omy27* we found genes related with innate immune response regulation, *NF-kB* activation by *TNFα*, and some molecules related with metabolic process and apoptosis. However, the SNP explaining the highest proportion of genetic variance is located within an exon of the gene *Smoothelin protein 2* (*Smtnl2*) which remains poorly characterized both in humans and fishes, but it is believed that participates in actin cytoskeleton organization.

The complete list of genes located within the 1Mb window flanking the SNPs explaining the highest proportion of genetic variance, within each genomic region associated with resistance to *P. salmonis*, is shown in Table S1.

## DISCUSSION

In the current study we show significant genetic variation for resistance to *P. salmonis* in a farmed rainbow trout population. A moderate to high heritability was estimated for resistance as TD (0.48) and BS (0.34). These estimates are higher than those reported in previous studies carried out for resistance to other bacterial diseases in aquaculture species, with heritabilities ranging from 0.22 to 0.38 (Ødegård *et al.* 2006; Palaiokostas *et al.* 2016; Vallejo, *et al.* 2017b). In the case of *P. salmonis* resistance, several studies have evaluated the presence of genetic variation in different salmonid species. Thus, similar estimates have been shown for Atlantic salmon, when using pedigree or genomic data, with values ranging from 0.19 to 0.39 (Yáñez *et al.* 2013; Yáñez, *et al.* 2014b; Correa *et al.* 2015; Bangera *et al.* 2017). In the case of coho salmon, heritability estimates range from 0.16 to 0.27 when resistance is defined as a linear or binary trait (Yáñez, *et al.* 2016a; Barría, *et al.* 2018).

Recent studies in rainbow trout, using different pedigree and genome-based genetic evaluation approaches, estimate heritabilities ranging from 0.39 to 0.57 for TD and from 0.54 to 0.62 for BS (Yoshida, *et al.* 2018a); values which are within the range of our estimations. Moreover, our results suggest a higher effect of the additive genetic component on the phenotypic variance for resistance to *P. salmonis* in rainbow trout when compared to *S. salar* and *O. kisutch*, which would imply potentially faster genetic progress for the improvement of resistance to *P. salmonis* by means of artificial selection in the rainbow trout population used in the present study.

The effect of the genetic architecture of a trait (among other variables) on the accuracy of breeding values obtained through genomic selection (GS) is widely known (Daetwyler *et al.* 2008; Goddard 2009). Previous studies in salmonid species (Atlantic salmon and coho salmon), suggest that resistance to *P. salmonis* is a polygenic trait (Correa *et al.* 2015; Barría, *et al.* 2018). Based on the 26K SNPs which passed QC, our study similarly suggests a polygenic nature for resistance to *P. salmonis* resistance in rainbow trout (*i.e.* no QTL explaining >= 10% of the genetic variance). Thus, it is expected that, when compared with a pedigree-based Best Linear Unbiased Predictor (BLUP) method, a genomic BLUP approach for GS would have an increase in accuracy of breeding values over a Bayesian approach (Habier *et al.* 2007; Hayes *et al.* 2009) for the current rainbow trout population. Nonetheless, as predicted by Yoshida, *et al.* (2018b) this was true only at low SNP densities (*i.e.* 0.5 to 10 K). When 20K and 27K were used, Bayes C outperformed GBLUP accuracies. The authors suggested that this could be due to an oligogenic architecture of the resistance trait, or that Bayes C had higher effectiveness in capturing the linkage disequilibrium between the SNPs and a QTL when more SNPs were used.

Resistance to bacterial infections implies a modulation of the host immune response to inhibit or reduce the replication rate of the pathogen (Doeschl-Wilson and Kyriazakis 2012). The infection process carried out by *P. salmonis* uses clathrin for internalization and then the actin cytoskeleton for vacuole generation (Ramírez *et al.* 2015). Similar pathways have been observed in other mammalian intracellular gram-negative bacteria (Manon *et al.* 2012; Valencia-gallardo *et al.* 2015). Within the region associated with TD on *Omy3* we identified a gene coding for the receptor DC-SIGN related with the immune response and expressed on macrophage and dendritic-cell surfaces (Ahmed *et al.* 2015). It has been previously described that *Mycobacterium tuberculosis*, interferes with the Toll-like receptor signaling by DC-SIGN, inhibiting interleukin-12 production (Gorvel *et al.* 2014), a proinflammatory cytokine, which plays a key role in the performance of phagocytes in teleost fish (Alvarez *et al.* 2016).

As mentioned before, endocytosis mediated by clathrin is the main pathway used by *P. salmonis* for cell invasion. Clathrin recruits, among other cell components, AP-2; which is regulated by NECAP-1 (Ritter *et al.* 2013), a gene flanking the SNP explaining the highest proportion of genetic variance in *Omy3* for resistance measured as TD. Similarly, on this chromosome we also found the gene *glutathione S-transferase kappa 1* (*gstk1*) (GTS), which is member of the glutathione S-transferase family (GST), involved in cellular detoxification, and expressed in cells to reduce oxidative stress-related damage (Morel and Aninat 2011), a consequence of *P. salmonis* infection (Rozas and Enríquez 2014), and differentially expressed in Atlantic salmon after *P. salmonis* exposure (Rise *et al.* 2004). A candidate gene related to resistance as measured by BS, was found on *Omy5*, the *fas ligand* gene (*faslg*) is a member of the TNF superfamily. The Fas/FasL pathway is essential for immune system regulation, including apoptosis induced by T cell activation and by cytotoxic T lymphocytes (Siegel *et al.* 2000).

For both resistance trait definitions, the same chromosome and identical SNP was identified as the marker explaining the highest genetic variation for resistance, which makes this QTL as an interesting region in rainbow trout. Within this region we found the gene *phosphatidylinositol transfer protein alpha* (*pitpna*), which belongs to the phosphatidylinositol family (ptdlns) (Piscatelli *et al.* 2016), and is responsible for phospholipid transfer between cellular membranes (Thornbrough *et al.* 2016), which in turn are regulators of cell signal transduction, membrane trafficking and cytoskeleton organization (Hilbi and Haas 2012). The latter process is affected by *P. salmonis* once inside the macrophages (Ramírez *et al.* 2015). Similar to *P. salmonis*, *Legionella pneumophila* also replicates inside macrophages, and manipulates the vesicle generation inside the cell by joining with ptdlns 5 (Hilbi and Haas 2012).

Additionally, in this region we found the gene *nlr family card domain containing 3* (*nlrc3*). Previously, Álvarez *et al.* (2017), described a higher differential expression of *nlrc3* in rainbow trout in response to bacterial lipopolysaccharides (lps), specifically in the skin, liver and gills. This pattern has also been observed in Atlantic salmon during an infection with *P. salmonis* (Tacchi *et al.* 2011), and is therefore a likely mechanism used by this bacteria to evade the immune response.

The gene *tapsain* (*tap*) is also involved in the immune response, transporting cytosolic peptides generated by the proteasome to load on MHC class I (Procko *et al.* 2005). On *Omy27*, we found a gene that encodes a protein related to tapsain (TAPBPR), which negatively regulates *tap*; generating a reduction in immune response efficiency (Boyle *et al.* 2013).

We expect that in the near future, the identification and validation of causative mutations affecting some of the candidate genes presented here, by means of functional studies, will provide a better understanding of resistance against this and other infectious diseases in rainbow trout and other salmonid species. These studies will be facilitated through international collaborative initiatives such as the Functional Annotation of All Salmonid Genomes, FAASG (Macqueen *et al.* 2017).

## CONCLUSIONS

To the best of our knowledge this is the first report identifying candidate genes related to resistance to *P. salmonis* in a farmed rainbow trout population. Genes likely related with resistance were identified close to SNPs explaining the highest proportion of genetic variance. Furthermore, we identified the same genomic region associated with resistance using both a linear and binary trait. Our results show that this trait is controlled by multiple genes each with a small effect. Therefore, a genomic selection approach is suggested as the best method to improve this trait by means of artificial selection.

## Supporting information

10.6084/m9.figshare.7883342

## ACKNOWLEDGMENTS

We would like to thank Aguas Claras S.A. for providing funding for the experimental challenge and fish used in this study. This work was also partially funded by the grant CORFO Innova-Chile (11IEI-12843) and FONDEF NEWTON-PICARTE (IT14I10100), funded by CONICYT (Government of Chile) and the Newton Fund - The British Council (Government of United Kingdom). JMY is supported by Núcleo Milenio INVASAL funded by Chile’s government program, Iniciativa Científica Milenio from Ministerio de Economía, Fomento y Turismo.

## AUTHORS’ CONTRBUTIONS

RM-N assessed the GWAS analyses, genes identification and contributed with discussion. AB wrote the initial version of the manuscript and contributed with discussion. PC contributed with discussion. MEL contributed with initial analysis. LB performed DNA extraction. JPL contributed with study design. JMY conceived and designed the study and supervised the work of RM-N. All authors reviewed and approved the manuscript.

### Animal ethics approval

Experimental challenge was approved by the Comité de Bioética Animal from University of Chile (Certificate Number 17041-VET-UCH)

