## Supplementary material for "Single-step genome-wide association study for resistance to *Piscirickettsia salmonis* in rainbow trout (*Oncorhynchus mykiss*)": 10.6084/m9.figshare.7883342

**Table S1.** Full list of genes located within top ten 1-Mb windows associated with P. salmonis resistance in rainbow trout, for TD and BS.

| CHR - SNP | Genes |
| --- | --- |
| Resistance as Time to Death (TD) | |
| 27 - Affx-88923370 | *CAPNS1-like, SNAP25-like, Unc-LOC11050749, RAB1A-like, TIMM50-like, Unc-LOC110507498, DLC, CHP2-like, RNASET2-like, SUPT5H, Unc-LOC110507504, PLEKHG3-like, SPINT2-like, PPP1R14A-like, TNFAIP2-like, TBCEL-like, GIMAP8-like, FAM57A-like, GEMIN4-like, TAPBPL-like, SMTNL2, POLDIP2-like, Unc-LOC110507518, SEBOX-like, CQ032, LRRC75A-like, CRK-like, YWHAE, MYO1C-like, NLRC3-like, SLC43A2-like, PITPNA-like, INPP5K-like, TEKT1, FBXO39-like, XAF1-like, RAP1GAP2, Unc-LOC110507536, BLMH, SLC6A4-like, EFCAB5, SSH2-like.* |
| 3 - Affx-88916453 | *GRIN2D-like, MYOCD-like, Unc-LOC110508932, CSL2-like, Unc-LOC110508936, STL2, PITPNC1-like, Unc-Mitochondrial protein AtMg00860-like, SRRM2-like, CACNG7-like, Unc-LOC110508944, CACNG6-like, U2AF2-like, CCDC106-like, TMEM238-like, SPAG9-like, RCVRN-like, GSG1L, MFAP5-like, NECAP1-like, AICDA-like, NAT14-like, ZNF729-like, IL11-like, RASIP1-like, COX6B1, SLC2A1-like, TRIM21-like, EPHB5-like, TRPV6-like, Unc-LOC110505091, TRPV5-like, CD209E, GABARAPL1-like, ARHGEF5-like, Unc-LOC110520910, M6PR-like, PHC1-like, STYK1-like, GSTK1-like, RAP1GAP2-like, CASP2-like, Unc-LOC110509290, PERK10-like, Unc-LOC110509304, LOC110520913, KEL-like, EMG1-like, HIST1H1-like, NSUN5, IGFBP4-like, NCAPD2.* |
| 14 - Affx-88922612 | *GJB1-like, SPINK1-like, CREB5-like, TLR4-like, OSBPL3-like, NSHIP5-like, MPP6-like, CREB5 (pseudogene), JAZF1, TAX1BP1, NPY, CCDC126-like, TRA2A-like, IGF2BP3-like, QNR71-like, ZG57-like, OXNAD1-like, Unc-OC110487979, DAZL, PLCL2, TBC1D5, SATB1-like, KCNH8-like, RAB5A-like, KAT2B-like, ZNF385D-like, RAB25-like.* |
| 24 - Affx-88927397 | *MECOM-like, PNPLA8-like, Unc-LOC110503452, OVOS2-like, ACTN3-like, A2M-like, USF1-like, GLCAK1-like, UBASH3B-like, SORL-like, Unc-LOC110503764, BCO2-like, SDHD-like, IL4I1-like, SNX19, ADAMTS15-like, ADAMTS8-like, ZBTB44-like, C11ORF53, Unc-LOC110503774, POU2AF1, NFKBIZ-like, Unc-LOC110503777, Unc-LOC110503783,* |
| Resistance as a Binary Survival (BS) | |
| 27 - Affx-88923370 | *CAPNS1-like, SNAP25-like, Unc-LOC11050749, RAB1A-like, TIMM50-like, Unc-LOC110507498, DLC, CHP2-like, RNASET2-like, SUPT5H, Unc-LOC110507504, PLEKHG3-like, SPINT2-like, PPP1R14A-like, TNFAIP2-like, TBCEL-like, GIMAP8-like, FAM57A-like, GEMIN4-like, TAPBPL-like, SMTNL2, POLDIP2-like, Unc-LOC110507518, SEBOX-like, CQ032, LRRC75A-like, CRK-like, YWHAE, MYO1C-like, NLRC3-like, SLC43A2-like, PITPNA-like, INPP5K-like, TEKT1, FBXO39-like, XAF1-like, RAP1GAP2, Unc-LOC110507536, BLMH, SLC6A4-like, EFCAB5, SSH2-like.* |
| 5 - Affx-88951679 | *RNF170-like, PFAS, IPO13-like, PERK2-like, Unc-LOC110524487, MMACHC, LRRC52-like, Unc-LOC110524491, ZYG11B, COA7, RALGPS2, ANGPTL1-like, FAM20B, FASLG-like, Unc-LOC110524499, JUN, PRDX6-like, PLPP6, FMO1, Unc-LOC110524500, FMO5-like, PRRC2C-like, VAMP4-like, MYOC-like, METTL13-like, ITPA-like, DNM2-like, SUCO-like.* |
| 29 - Affx-88908715 | *NCOA3-like, RBM38-like, MIP-like, AQP4-like, BAZ2A-like, PTGES3-like, NACA, Unc-LOC110532028, Unc-LOC110532029, RHOD6-like, DDX23, CACNB3-like, SPRYD4-like, GLS-like, COL2A1-like, AAAS-like, ADCY6-like, Unc-LOC110532032, Unc-LOC110532033, G6PD, GPD1, ASIC1-like, MAPK12-like, PCBP2, KIF5B-like, Unc-LOC110532046, MBD6-like, STAC3-like, NXPH4-like, NDUFA4-like, ACTR5, SLC32A1-like, PRELID3B-like, ATP5F1E-like, PIEZO2-like, GNB1-like, VAPB-like, CHMP4C, EIF2S2, RAE1, SRSF6-like, L3MBTL1-like, SGK2-like.* |

Unc. – Uncharacterized
